# Robust Determination of Protein Allosteric Signaling Pathways

**DOI:** 10.1101/565457

**Authors:** Wesley M. Botello-Smith, Yun Luo

## Abstract

To understand how protein function changes upon an allosteric perturbation, such as ligand binding and mutation, significant progress in characterizing allosteric network from molecular dynamics (MD) simulations has been made. However, determining which amino acid(s) play an essential role in the propagation of signals may prove challenging, even when the location of the source and sink is known for a protein or protein complex. This challenge is mainly due to the large fluctuations in protein dynamics that cause instability of the network topology within a single trajectory or between multiple replicas. To solve this problem, we introduce the current-flow betweenness scheme, originated from electrical network theory, to protein dynamical network analysis. To demonstrate the benefit of this new method, we chose a prototypic allosteric enzyme (IGPS or HisH-HisF dimer) as our benchmark system. Using multiple replicas of simulations and multiple network topology comparison metrics (edge ranking, path length, and node frequency), we show that the current-flow betweenness provides a significant improvement in the convergence of the allosteric networks. The improved stability of the network topology allows us to generate a delta-network between the apo and holo forms of the protein. We illustrated that the delta-network is a more rigorous way to capture the subtle changes in the networks that would otherwise be neglected by comparing node usage frequencies alone. We have also investigated the use of a linear smoothing function to improve the stability of the contact map. The methodology presented here is general and may be applied to other topology and weighting schemes. We thus conclude that, for determining protein signaling pathways between the pair(s) of source and sink, multiple MD simulation replicas are necessary and the current-flow betweenness scheme introduced here provides a more robust approach than the geodesic scheme based on correlation edge weighting.

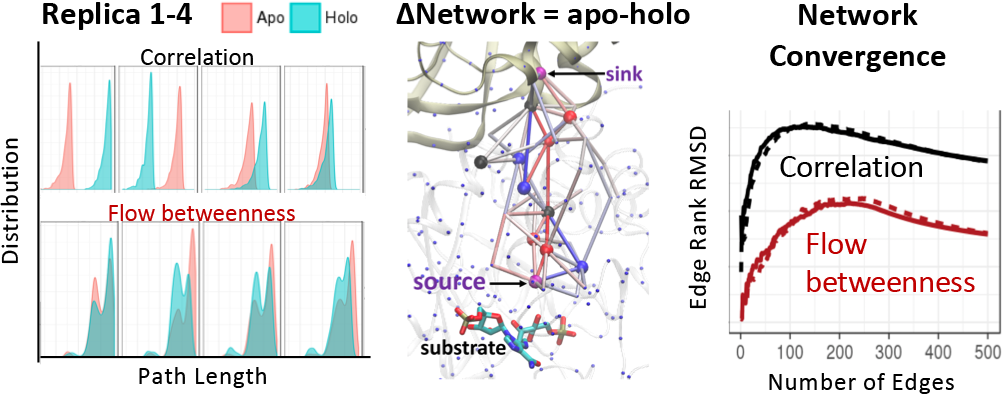
For Table of Contents Only

## Introduction

Allosteric regulation is ubiquitous inside the cell and serves to regulate cell functions in a delicate way. To understand how protein function changes upon external perturbation, such as protein-protein/ligand binding, mutation, or voltage change, numerous researchers have implemented variants of allosteric network concepts to biomolecules^1-12^. One important application of network theory in the context of structural dynamics of biomolecules is to determine which amino acid residues in a protein play an essential role in the propagation of signals within a protein or between proteins by constructing a network of interactions between amino acid residues. Various computational approaches in identifying such allosteric pathways have been extensively reviewed^13-16^. Among them, graph theory based network analysis has been seeing increasing attention over the past few decades.

In graph theory, a network topology is usually defined by a set of nodes representing structural features such as the alpha carbon or side chain of amino acids, and a set of edges connecting the nodes, representing interactions between these features. To assess the importance of a given node or edge with respect to signal propagation, common methods make use of betweenness centrality, also known as geodesic centrality, which measures the number of shortest paths that cross a given node or utilize a given edge in the studied network. This measurement is quite straightforward. It provides a useful metric to determine which nodes are important with respect to general propagation of signals (e.g., energy, motion, conformational change, etc.) across the network.

While betweenness centrality is useful in the case of an exploratory study, often, one is concerned with the propagation of changes between a known pair or set of residues or domains. For example, how a ligand binding or a single mutation on a protein allosterically alters the dynamics of a remote site on the same protein or protein oligomers. When applied under that context, betweenness centrality may contain contributions from paths that are not relevant to transmitting a signal between the domains of interest. On a related note, betweenness centrality considers only the shortest paths between nodes in a network. However, edges and nodes that lie ‘near to’ but not exactly on the shortest path that may provide relevant contributions will be ignored by a standard betweenness centrality computation. These two drawbacks can lead to artifacts such as undervaluing the importance of nodes and edges that are near but are not part of shortest paths, and also to overvaluing of edges that lie on non-relevant paths. Moreover, since edges lying near to shortest paths may never be included at all, this can lead to significant instability when applying betweenness centrality to molecular dynamics simulations, wherein the network topology may fluctuate over time. The instability of network topology has been discussed previously^6^ and thoroughly demonstrated in the current study.

When the transmission between a pair of known residues or domains is of interest, a modified approach to betweenness centrality may be employed. First, one computes the shortest paths connecting residues in the two domains of interest. Next, contributions of edges and nodes that lie ‘near to’ but not on the shortest path may be included by assigning a cutoff value for path length. This method called “suboptimal path search” was developed and implemented in the *subopt* program from the NetworkView plugin of VMD program by Luthey-Schulten *et. al.* a decade ago ^2, 17-18^. An enhanced version of the *subopt* program called ‘Weighted Implementation of Suboptimal Paths’ (WISP)^7^ was recently developed, which achieves rapid calculation of suboptimal pathways, and may compensate, at least in part, for these fluctuations in the network topology by choosing a sufficiently large path length cutoff.

While the suboptimal path approach addresses the main shortcomings of the general geodesic betweenness metric described above, it is often difficult to know a priori the appropriate cutoff value for additional path length, which is obviously system dependent. In addition, computing sets of suboptimal paths for extremely long cutoff lengths can become quite cumbersome. Here, we show that there is an alternative betweenness metric that is generalizable to any system, network topologies, and weighting schemes, with significantly improved stability. This metric, known as current-flow betweenness, takes its roots in the analysis of electrical circuits^19^. The applicability of electrical circuit analysis to thermodynamics problems was proposed by Oster *et. al.* in 1971, in which they stated that “most systems that can be analyzed with the network approach share one common property: the rate of energy transmission and a flow variable. In electrical networks these variables are voltage difference and current; in mechanics are force and velocity …”^20^. Interestingly, this method is analogous to a metric commonly employed in the analysis of information propagation networks, known as information centrality^21^, which was derived in a much different manner. It can be shown that information centrality and current-flow centrality yield essentially equivalent results mathematically^19^. Moreover, it has been shown that these metrics are upper or asymptotic limits for certain random walk based methods^19, 22^. Here, we will focus on the current-flow betweenness formalism since the derivation is conceptually simpler and more intuitive than the derivation of the analogous information centrality measurement.

The current-flow betweenness metric may be applied (and in fact is most easily applied) when one is concerned only with paths connecting a specific set of nodes. Thus it does not suffer from the drawback of standard betweenness centrality where paths connecting extraneous pairs of nodes may lead to overvaluing the importance of a node or edge. Like the suboptimal path search, current-flow betweenness includes contributions from edges and nodes that are ‘near to’ but do not lie on a shortest path. However, the unique advantage of our method for searching suboptimal path lies in the fact that current-flow betweenness includes contributions from all possible paths connecting the set of nodes of interest (source and sink), not just those falling within a specified cutoff. Thus, in some sense, current-flow betweenness proposed here can be conceived of as a “cutoff-free” and more robust version of the suboptimal path method with respect to computing betweenness.

In this work, we introduce, for the first time, the application of current-flow betweenness to investigate allosteric networks in proteins. We demonstrated the advantage of current flow betweenness over correlation weighted betweenness in the scenario of ligand-binding induced allosteric network change. Our benchmark system is imidazole glycerol phosphate synthase (IGPS or HisH-HisF dimer), a prototypic system most frequently used for network analysis^6-7, 23-25^. IGPS belongs to the glutamate amidotransferase family that regulates the histidine biosynthetic pathway. The catalytic triad is located at HisH domain, however, the binding of an allosteric effector PRFAR to the HisF domain increases the catalytic turnover number by thousands fold. The allosteric mechanism of IGPS has been extensively studied, both experimentally and computationally, and hence provided us with an ideal benchmark system for implementing the current-flow scheme.

Using multiple replicas of MD simulations and various network topology metrics (edge ranking, path length, and node frequency), we show that a current-flow betweenness scheme vastly improved the convergence of allosteric network topology. The improved stability of topology allows us to generate a delta-network between the apo and holo IGPS systems. We show that delta-network is a more rigorous way to capture the subtle change in the network than comparing node frequencies alone. We have also investigated the use of a linear smoothing function to generate a contact map, which has been suggested as a means of improving network topology stability. A set of python scripts for calculating current-flow betweenness scores is available to download at https://github.com/LynaLuo-Lab/network_analysis_scripts, which can be easily incorporated to use together with NetworkView tool in VMD program.

## Theory and Methods

### Current-flow betweenness

Current-flow betweenness is framed in terms of a network of electrical resistor connecting a pair (or set) of source nodes and sink nodes across which a sufficient voltage is applied to induce a unit of current to flow. In the case of a system of electrical resistors, it is known that current will flow through all possible paths between a given source and ground simultaneously. If one were to measure the current flowing through a given edge (resistor) in such a network, this would then yield the “current-flow betweenness”, i.e. as the current-flow through that node or edge in the electrical network. For simple networks, this may be computed quite easily by hand by employing Kirchhoff’s laws. Figure 1 shows an example of using Kirchhoff’s Laws to compute current-flow through each edge of a network of resistors. As mentioned earlier, it can be shown that this ‘current-flow betweenness’ is equivalent to information centrality and several betweenness scoring metrics based on random walk approaches. These metrics have been shown to provide better results as compared with shortest path betweenness in other network analysis fields. A more detailed in depth derivation of flow betweenness can be found in the paper by Brandes and Daniel^19^, along with discussions of how it can be shown to be equivalent to information centrality.

**Figure 1:**
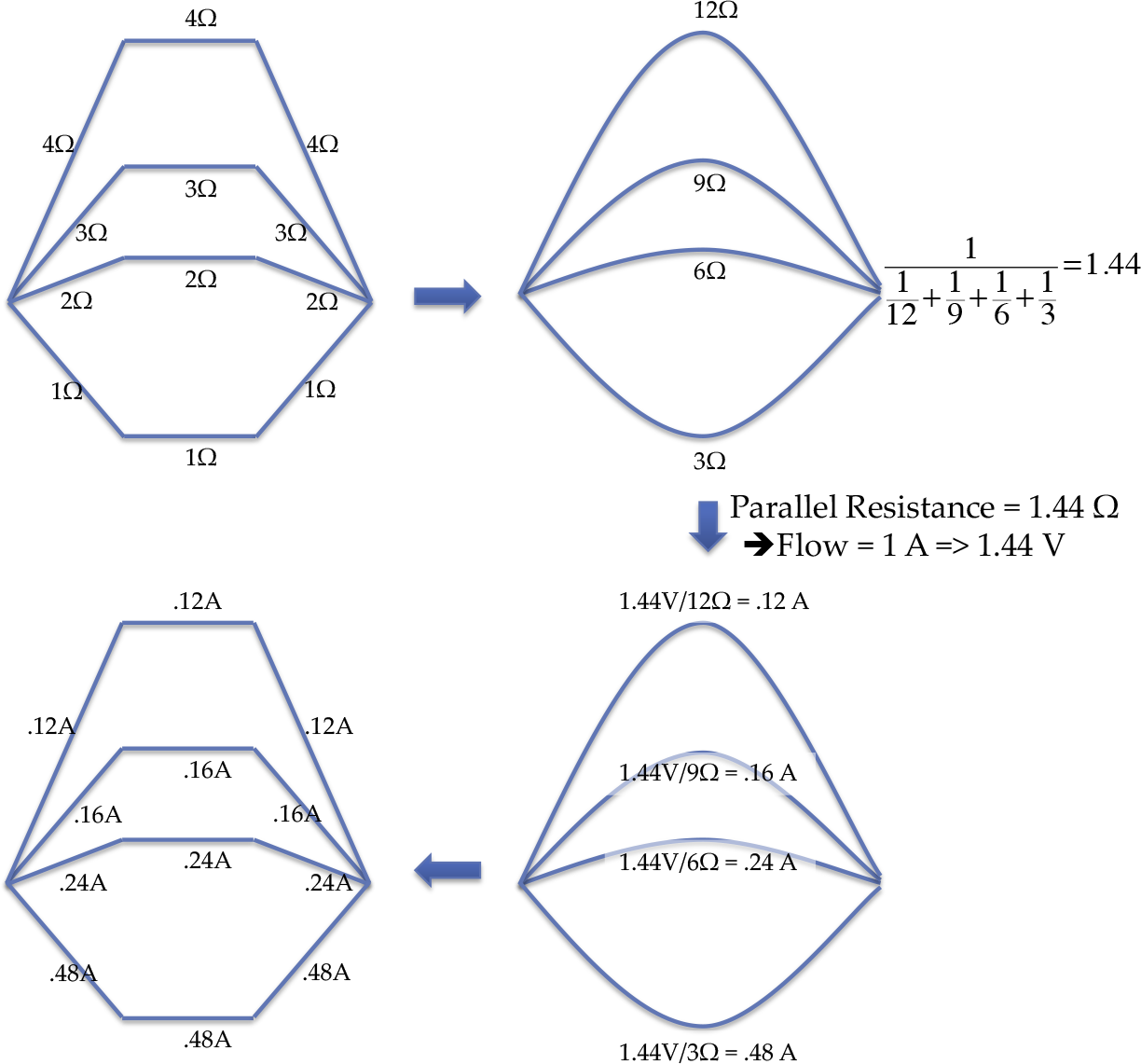
Example of using Kirchhoff’s Laws to compute current-flow through each edge of network of resistors.

To compute current-flow betweenness, one must first construct the associated adjacency and Laplacian matrices. The adjacency matrix, **A**, has as its elements, **a_i,j_**, which is the absolute values of the corresponding network weights **w_i,j_**. In the case of a network of electrical resistors, these weights would correspond to the conductance of each resistor. Here, we are interested in transfer of motion as described by a contact map weighted by correlation of atomic motions. Thus the absolute values of the entries in the correlation matrix will serve as the weights. I.e. given the atomic correlation matrix **C**, we assign **a_i,j_ = |c_i,j_|** if i, j corresponds to an edge in the contact map, and we assign **a_i,j_ =** 0 if i,j edge is not included in the contact map. The most frequently used criteria for contact map generation is: residue i is determined to be in contact with another residue j, if for at least 75% of the frames in a given trajectory, there is at least one atom of i in contact (e.g. within 4 Å) with at least one atom in j. We later showed that a smoothed function can be applied to increase the stability of the contact map (Table 1). **c**_i,j_ can be any correlation coefficients such as the commonly used Pearson correlation coefficients, or generalized correlation coefficients based on the mutual information concept^26^, or a more protein specific inter-residue interaction energy based weights^6^.

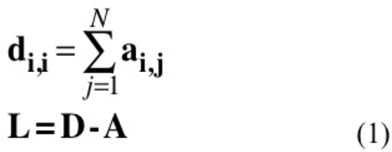

The difference between the diagonal matrix **D** and the adjacency matrix **A** is known as the Laplacian matrix **L** (Eq. 1), in which **d_i,i_** are the elements of the diagonal matrix **D** (the sum of the weights of edges connected to node i), constructed by summing over each row of the adjacency matrix **A**. Next, the inverse of this network Laplacian **L** must be computed to obtain the betweenness of each edge, as illustrated in Figure 2. Betweenness of each node can be computed by summing the betweenness of each connected edge. However, by its construction, the Laplacian matrix is guaranteed to be ‘singular’, meaning that no unique inverse exists. Fortunately, an appropriate ‘pseudo-inverse’ can serve just as well. The Moore-Penrose pseudo inverse is used here, although other methods such as single value decomposition may also work. It should be noted that computation of matrix inverses and pseudo-inverses can become quite computationally taxing for very large networks. However, the process is reasonably fast for networks consisting of thousands or even tens of thousands of nodes. In the case of allosteric network applications, network nodes are typically taken to represent individual residues within a protein and are computed from molecular dynamics or other simulation techniques that have similar size limitations and so, direct computation of pseudo-inverses should be tractable. In the event where they are not, approximation methods are available, such as the eigenvalue decomposition methods given in the paper by Bozzo and Franceschet^27^.

**Table 1:**
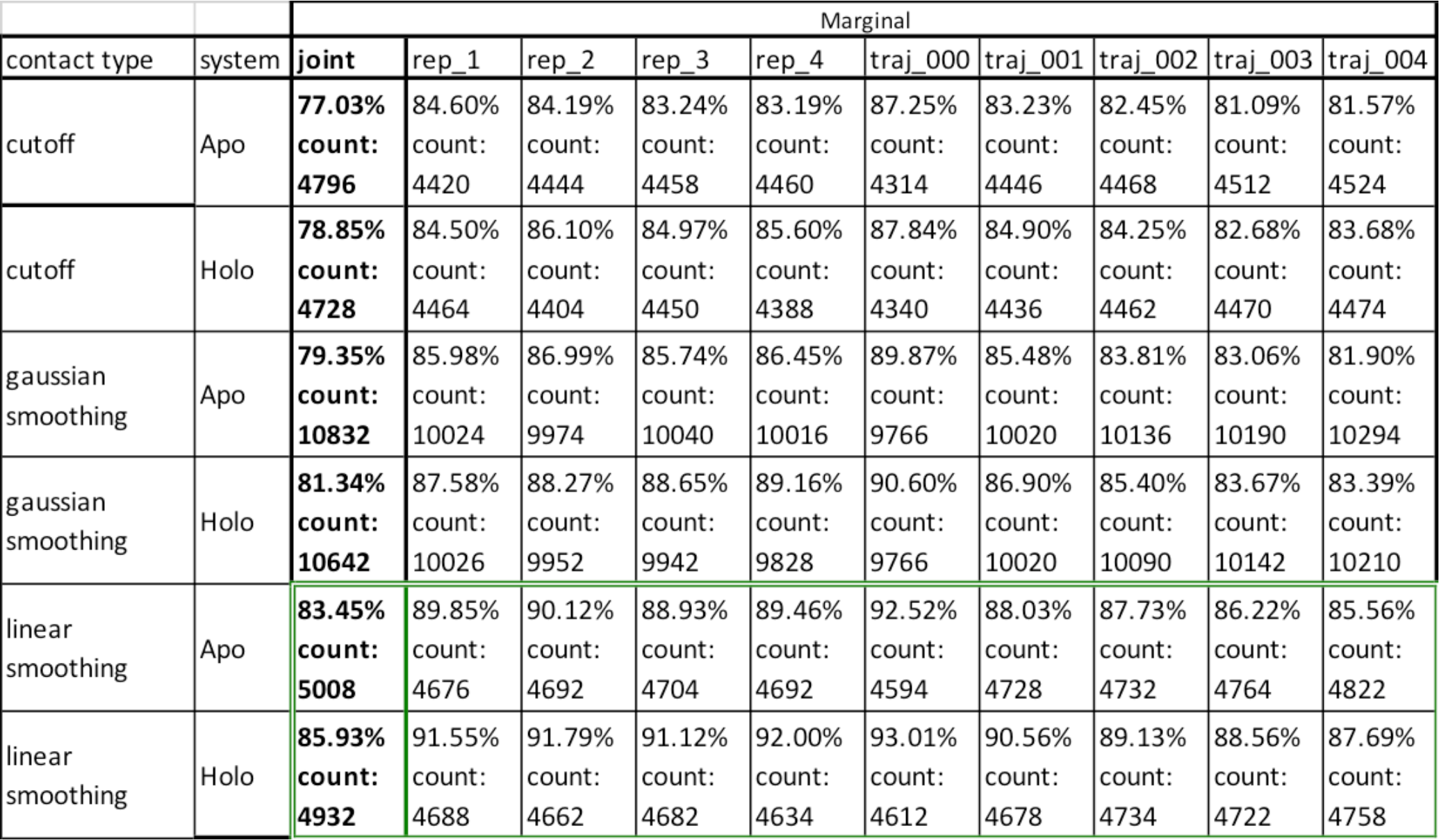
Marginal and joint stabilities of the contact map calculated for each system and each smoothing type. Count is the number of contact in the contact matrix.

| contact type | system | Marginal |  |  |  |  |  |  |  |  |  |
| --- | --- | --- | --- | --- | --- | --- | --- | --- | --- | --- | --- |
|  |  | joint | rep_1 | rep_2 | rep_3 | rep_4 | traj_000 | traj_001 | traj_002 | traj_003 | traj_004 |
| cutoff | Apo | **77.03%**<br>**count:**<br>**4796** | 84.60%<br>count:<br>4420 | 84.19%<br>count:<br>4444 | 83.24%<br>count:<br>4458 | 83.19%<br>count:<br>4460 | 87.25%<br>count:<br>4314 | 83.23%<br>count:<br>4446 | 82.45%<br>count:<br>4468 | 81.09%<br>count:<br>4512 | 81.57%<br>count:<br>4524 |
| cutoff | Holo | **78.85%**<br>**count:**<br>**4728** | 84.50%<br>count:<br>4464 | 86.10%<br>count:<br>4404 | 84.97%<br>count:<br>4450 | 85.60%<br>count:<br>4388 | 87.84%<br>count:<br>4340 | 84.90%<br>count:<br>4436 | 84.25%<br>count:<br>4462 | 82.68%<br>count:<br>4470 | 83.68%<br>count:<br>4474 |
| gaussian smoothing | Apo | **79.35%**<br>**count:**<br>**10832** | 85.98%<br>count:<br>10024 | 86.99%<br>count:<br>9974 | 85.74%<br>count:<br>10040 | 86.45%<br>count:<br>10016 | 89.87%<br>count:<br>9766 | 85.48%<br>count:<br>10020 | 83.81%<br>count:<br>10136 | 83.06%<br>count:<br>10190 | 81.90%<br>count:<br>10294 |
| gaussian smoothing | Holo | **81.34%**<br>**count:**<br>**10642** | 87.58%<br>count:<br>10026 | 88.27%<br>count:<br>9952 | 88.65%<br>count:<br>9942 | 89.16%<br>count:<br>9828 | 90.60%<br>count:<br>9766 | 86.90%<br>count:<br>10020 | 85.40%<br>count:<br>10090 | 83.67%<br>count:<br>10142 | 83.39%<br>count:<br>10210 |
| linear smoothing | Apo | **83.45%**<br>**count:**<br>**5008** | 89.85%<br>count:<br>4676 | 90.12%<br>count:<br>4692 | 88.93%<br>count:<br>4704 | 89.46%<br>count:<br>4692 | 92.52%<br>count:<br>4594 | 88.03%<br>count:<br>4728 | 87.73%<br>count:<br>4732 | 86.22%<br>count:<br>4764 | 85.56%<br>count:<br>4822 |
| linear smoothing | Holo | **85.93%**<br>**count:**<br>**4932** | 91.55%<br>count:<br>4688 | 91.79%<br>count:<br>4662 | 91.12%<br>count:<br>4682 | 92.00%<br>count:<br>4634 | 93.01%<br>count:<br>4612 | 90.56%<br>count:<br>4678 | 89.13%<br>count:<br>4734 | 88.56%<br>count:<br>4722 | 87.69%<br>count:<br>4758 |

**Figure 2:**
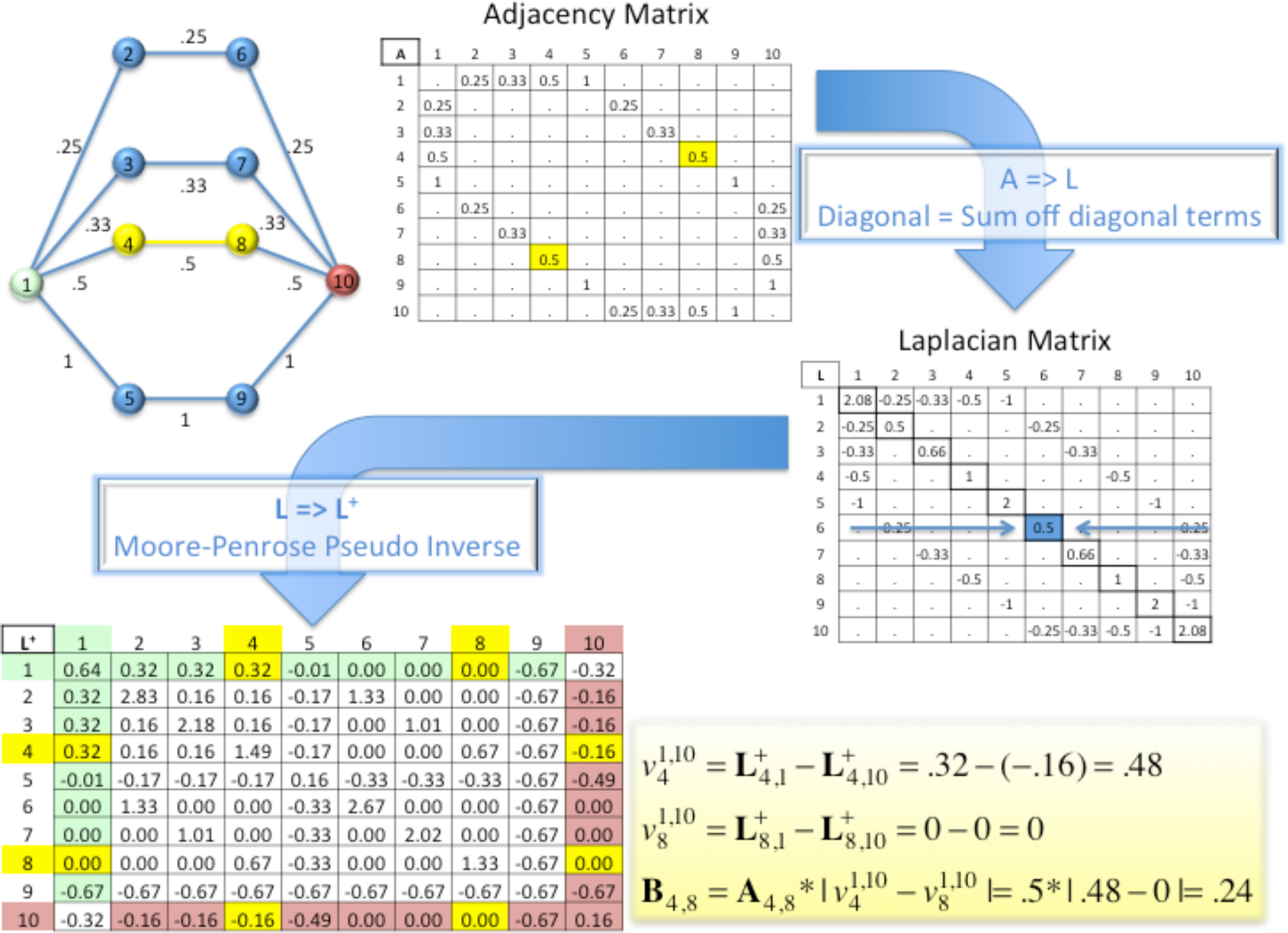
Calculation steps of flow betweenness for edge 4-8 of an example network (top left corner). Note that this network is equivalent to the network in Figure 1. Here, edges are labeled with conductance value rather than corresponding resistance values.

Given a network’s adjacency matrix, **A**, and a suitable inverse (or approximation thereof) to its Laplacian, **L^+^**, the current-flow betweenness between nodes ***i*** and ***j***, **B*_i,j_***, for a given source node ***s***, and target node ***t***, is given as in equation 2:

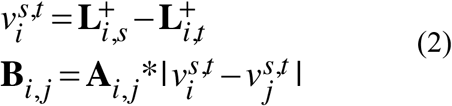

Where 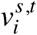 is the potential at a given node ***i***. In the case of multiple nodes as source and target, as in this study, one may attain the current-flow as the sum over all combinations of source and target nodes divided by the number of combinations. This is a relatively trivial double sum when the two sets are disjoint, e.g:

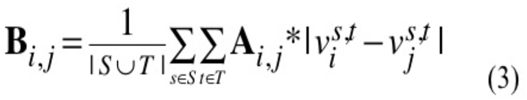

Where S, T are disjoint sets of source and target nodes respectively. When S and T are not disjoint (e.g. as in computing a generalized current betweenness over the entire network) one must take care to avoid / remove contributions from double summations^19^. Here we consider only the former case where the sources are disjoint from the targets. Figure 2 illustrates an example of flow betweenness calculation for a simple 10-node network with a single source and sink. The corresponding ‘by hand’ calculation using Kirchhoff’s Laws was shown previously in Figure 1 for the analogous network of electrical resistors, where each edge’s resistance is equal to the reciprocal of the correlation shown in Figure 2. One can see that the result for edge 4-8 (**B**_4,8_) using the above matrix formalism is identical to the result from electrical network analog in Figure 1.

### System Preparation and Simulation

We have simulated the IGPS systems in an equivalent manner to previous studies of the same system^6-7, 23^. Briefly, the crystal structure of HisH-HisF (PDB 1GPW) was used as substrate-free (apo) structure and the coordinate of PRFAR substrate was taken from the crystal structure (PDB 1OX5) to build substrate-bound (holo) state. CHARMM-GUI ^28^ was used to read in the PDB file and generate solvated systems. All simulations employed the all-atom CHARMM C36 force field for proteins and ions, and the CHARMM TIP3P force field^29^ for water. All molecular dynamics simulations were performed using the PMEMD module of the AMBER16 package^30^ with support for MPI multi-process control and GPU acceleration code. Orthorhombic periodic boundary conditions were used for all simulations in the isobaric-isothermal (NPT) ensemble using Langevin thermostat and Berendsen barostat. The pressure and temperature were maintained at 1 atm and 310.15 K. Long-range electrostatic interactions were treated using the default particle mesh Ewald (PME) method. Nonbonded cutoff is set to be 8 Å. The dynamics were propagated using Langevin dynamics with Langevin damping coefficient of 1 ps^−1^ and a time step of 2 fs. The SHAKE algorithm was applied to all hydrogen atoms. For each system (apo and holo), for four independent replicas (different initial velocities) of 50 ns production runs were accumulated for analysis. Protein snapshots were extracted at 100 ps intervals for calculating contact map and correlation coefficients.

### Dynamical Network Analysis

Amaro and colleagues examined various node representations in Cartesian coordinates, and reported that the amino acid residue center of mass (COM) performed better in detecting HisH:Lys181-HisF:Asp98 salt bridge known to participate in allostery^23^. We thus used pytraj to extract the residue COM trajectories from original all-atom trajectories, then use CARMA program to generate correlation matrices for residue COM or alpha carbon (CA). The resulting data matrices were then used to compute associated current-flow betweenness scores. To calculate suboptimal path, we chose the substrate binding residue HisF:Leu50 as the source node and HisH:Glu180 in the catalytic triad as the sink node, same as the nodes used in previous work on suboptimal paths search^7^.

## Results and Discussion

### Increased stability of contact map topology using linear smoothing function

In current dynamical network analysis, optimal and suboptimal pathways are generated based on a contact map. Residues within a particular distance of another residue for certain percentage of the simulation time are assumed to influence the communication pathway directly, and residue that do not satisfy these criteria are removed from analysis. Therefore, the fluctuation in the contact map has a direct consequence in the instability of the network topology. To reduce the fluctuation of the contact map, we tested three cutoff criteria: 1) The default contact criteria in the program CARMA^31^ (4.5 Å distance for at least 75% of an MD trajectory); 2) Linear smoothing from 3 Å full contact to 6 Å zero contact (‘half’ contact at 4.5 angstroms); 3) Gaussian smoothing with 3 Å full contact, ‘half’ contact at 4.5 Å, and σ =1 Å.

To measure the stability of the contact map, four replicas (rep_1 to rep_4) of 50 ns trajectory were split into five 10 ns windows (traj_001 to traj_004) to allow statistical analysis (Table 1). The stability is calculated by 1–normalized standard deviation of the contact frequency over the given marginal (i.e. all traj_001 over four replicas, or traj_001 to traj_004 over a single replica). The joint stability is the stability over all trajectories and all replicas. Since contact is measured as a binomial variable, i.e. either 1 if a residue pair is in contact or 0 if not, its standard deviation of contact *i* can be given in terms of the contact frequency by the equation:

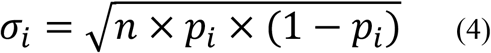

Where *n* is the number of frames over a marginal or joint set of trajectory windows, and *p*_*i*_ is the contact frequency of contact *i*. This, then, implies that the maximum possible standard deviation occurs when the contact frequency (*p*) is equal to 50%. Therefore, we may compute normalized standard deviations of contact frequency by dividing by 0.5√(n). Since standard deviation is a measure of how frequently a contact is formed or lost, we then define stability *s*=1-〈σ_*i*_〉. These results are tabulated in Table 1.

From table 1, we found that within the same system and same cutoff criteria, the stability of the contact map is similar between replicas and does not deviate alone simulation time (from traj_000 to traj_004). Comparing with the default cutoff criteria, Gaussian smoothing function yield larger number of contacts (counts), but only slightly improved stability, while the linear smoothing function provided consistent higher stability across all replicas, trajectories, and systems (highlighted in green box). It is of course possible to further fine tune the Gaussian smoothing function or introduce other types of cutoff to further improve the contact map stability. Our current data demonstrated that the linear smoothing function used here is superior to the default cutoff method.

### Significant improvement in the edge ranking convergence using current-flow betweenness

As discussed previously,^6, 8^ choice of weighting scheme can impact the network topology. We follow a similar protocol here to compare the stability of the network generated using current-flow betweenness weights versus correlation weights. The weight of each edge are either the -ln(|betweenness|) or -ln(|correlation|). Network occurrence frequency (*η*) is assigned to each edge by counting the number of paths the edge occurred over a total number of sub-optimal paths ranked by their flow betweenness weights or correlation weights.

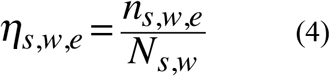

Where *e* is an edge in window *w*, of the system *s*, being considered. *n* is the number of paths that edge *e* occurs in and *N* is the total number of paths computed for window *w* of system *s*. These values were then used to rank each edge in each window with a rank of 1 assigned to the edges with the highest *η* value, rank of 2 to the next highest *η* value, etc. In practice, *subopt* program in VMD was run iteratively until at least *N* paths were generated and then only paths with rank less than or equal to *N* were taken. Since *subopt* program returns path lengths with only integer precision, ties can occur frequently. To handle ties, a maximum ranking scheme was employed in which all ties are assigned a rank equal to the next lowest rank plus the number of ties. Due to this method, replicas may contain slightly fewer or slightly more than *N* paths in cases where there were ties at ranks. Next, the average, <r(*η*)>_s,e_ was computed over all 20 windows for each system, followed by a second round of ranking based on each edge’s average *η* ranking R(<r(*η*)>)_s,e_. Finally the average root mean square deviation (RMSD) of edge ranks versus the number of edges included for calculating RMSD was plotted for each system to compare ranking based on correlation weights versus betweenness weights. E.g. the top *m* edges were selected based on average rank score and the RMSD of ranking over the 20 windows was computed using those top *m* edges.

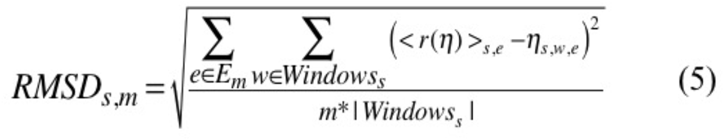

Where, *s* is one of the systems being considered, *m* is the number of top ranked edges (by ranking of average *η* values, R(<r(*η*)>)_s,e_, over all windows for system *s* and edge *e*, being considered). *E*_*m*_, is the set of those top *m* edges for system *s*, and |windows_s_| is the number of windows for the system. The results are shown in Figure 3. The <*η*> over four replicas for the 365 edges exhibiting non-zero frequency in at least one of the four top 700 suboptimal path networks (apo correlation, apo betweenness, holo correlation, holo betweenness) are provided in the supporting data (Table S1).

**Figure 3:**
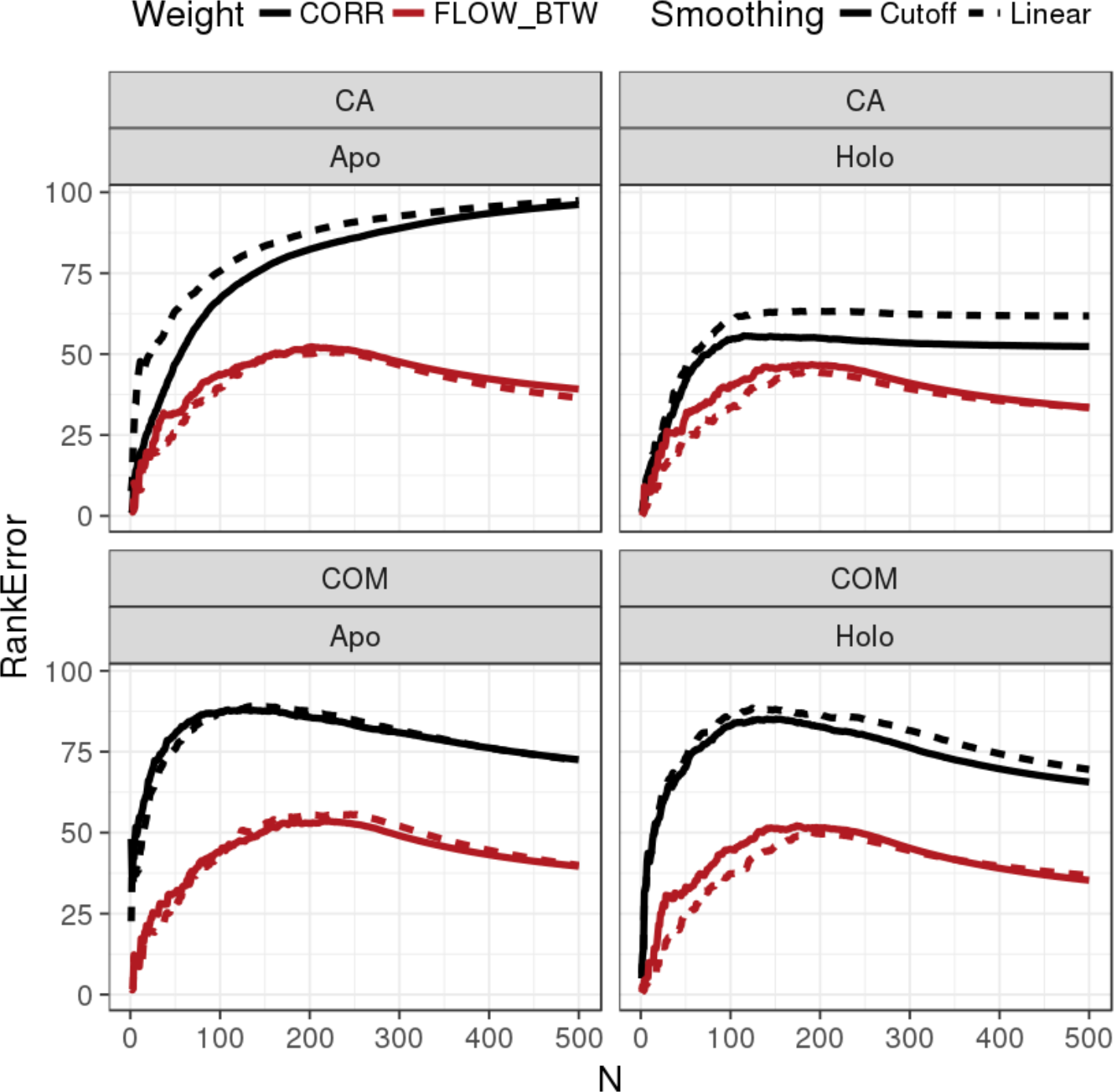
Comparison of edge ranking stability for correlation (black) and current flow betweenness (red) for apo and holo systems. For each system, the nodes representation is either alpha carbon (CA) or center of mass of each residue (COM). For each system, two different contact maps were used to generate either correlation based network (CORR) or current-flow betweenness (FLOW_BTW) based network: solid line represents the contact map generated using the default cutoff; dash line represents the linear smoothing cutoff contact map generated using linear smoothing function.

Figure 3 shows clearly that no matter which node representation or which contact map we used, for both apo and holo systems, flow betweenness weighting significantly decreases the edge ranking error (red line) compared with direct use of correlation as weights (black line), which means the network topology generated by flow betweenness scores is much more stable between simulation windows. The poor performance of using correlation to weight the edge is largely due to the fact that the change in correlation value of an edge only indicates the change of a specific edge, which may exhibit significant fluctuation. Furthermore, changes in the correlation between a pair of nodes may not necessarily imply an increase in path usage. This is particularly relevant when considering systems over which the residue-to-residue contacts and correlations may fluctuate significantly. It is again because current-flow betweenness score of an edge considers the contribution from all possible pathways between the source and sink nodes, and thus is more robust in capturing allosteric signaling transmission.

The linear smoothing function, which certainly improves the stability of the contact map, has negligible effect on the edge ranking error (Figure 3), suggest that the fluctuation of the network topology is largely due to the fluctuation in the edge weights during the simulations. Figure 4 illustrated two network topology of apo IGPS generated using the same contact map with edges colored based on their pairwise correlation or current-flow betweenness scores. The flow betweenness clearly show more focused signal transmission between source and sink, while the correlation contains no information of source and sink. It is worth noting that the flow betweenness metric can be applied to any types of contact map and weighting schemes, and thus can be combined with other methodology, such as energy-based weighting scheme^8^.

**Figure 4.**
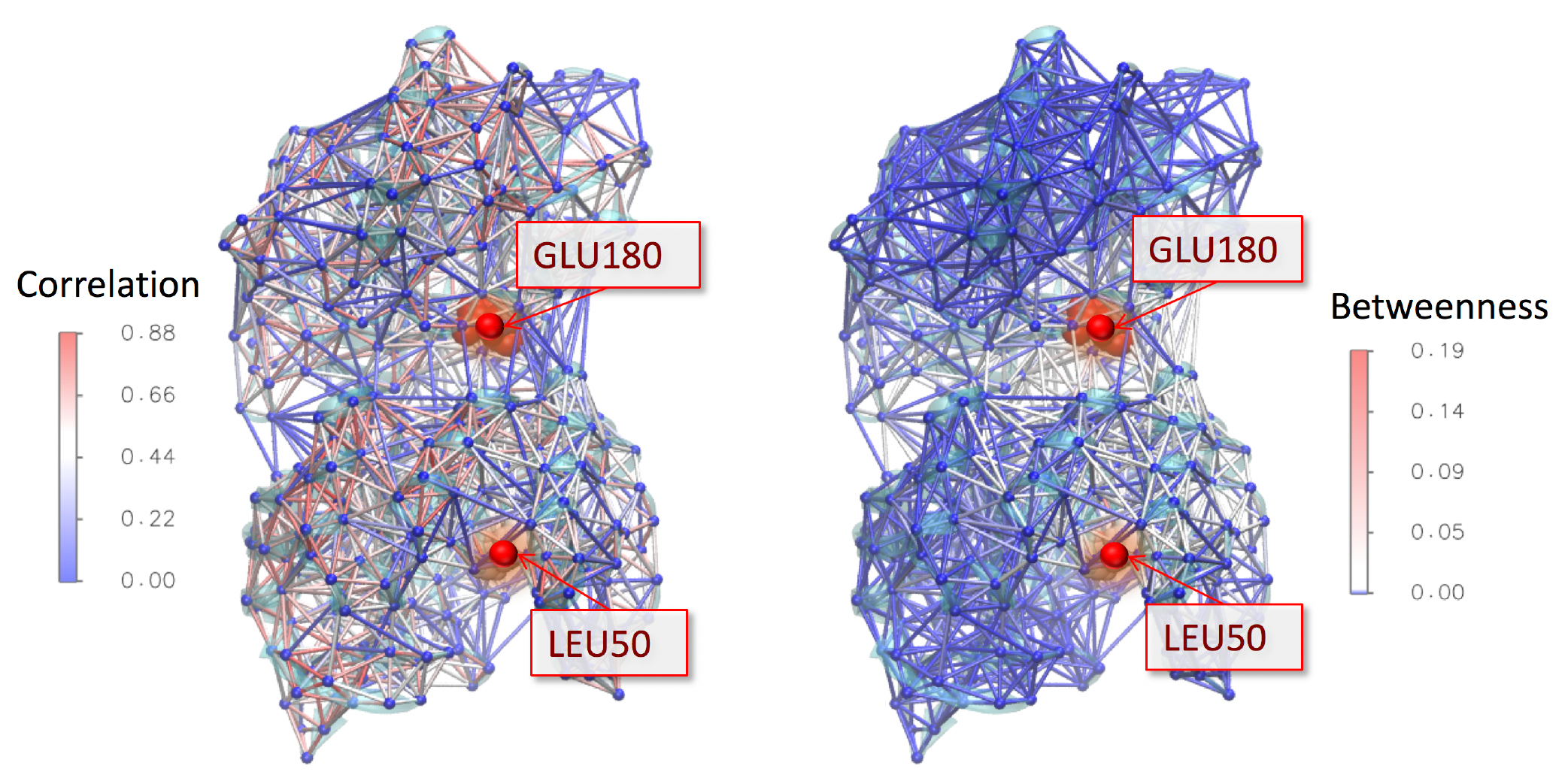
Projection of absolute Pearson correlation (left) and current-flow betweenness (right) edge weighted networks onto the structure for the apo IGPS protein backbone. Betweenness is calculated using GLU 180 and LEU 50 as source and sink nodes. Source and sink residues are highlighted in red VDW mode. Edges are colored based on their correlation score or betweenness score.

### Comparison of distribution of path length between replicas of simulations

Besides the edge ranking, the distribution of the path length among hundreds of suboptimal path is an important feature of the allosteric communication. For example, the allosteric communication may be altered by a shift in the path length distribution without altering the shortest path or the average suboptimal path length. We calculated the path length of the top 700 suboptimal paths ranked by their distance between source and sink for each replicas. The path length was calculated using either the sum of -ln(|correlation|) over the edges connecting the path or using the sum of -ln(|betweenness|). Our results (Figure 5) clearly show that when using the correlation edge weighting, the path length distribution is not consistent between replicas, especially between replica 1 and replica 2. When using current-flow betweenness edge weighting, the path length distributions are much more consistent among four replicas. All four replicas yield similar multimodal distribution in the apo and holo systems. While replica 1 showing minimum shift in the distribution, all other replicas consistently show shorter path length in the holo than apo network. This observation is in agreement with a previous allosteric network study suggesting that the motions of the residues connecting the source and sink sites are more tightly correlated when the substrate PRFAR bound to IGPS, possibly indicating loss of entropy along the allosteric pathways^7^.

**Figure 5.**
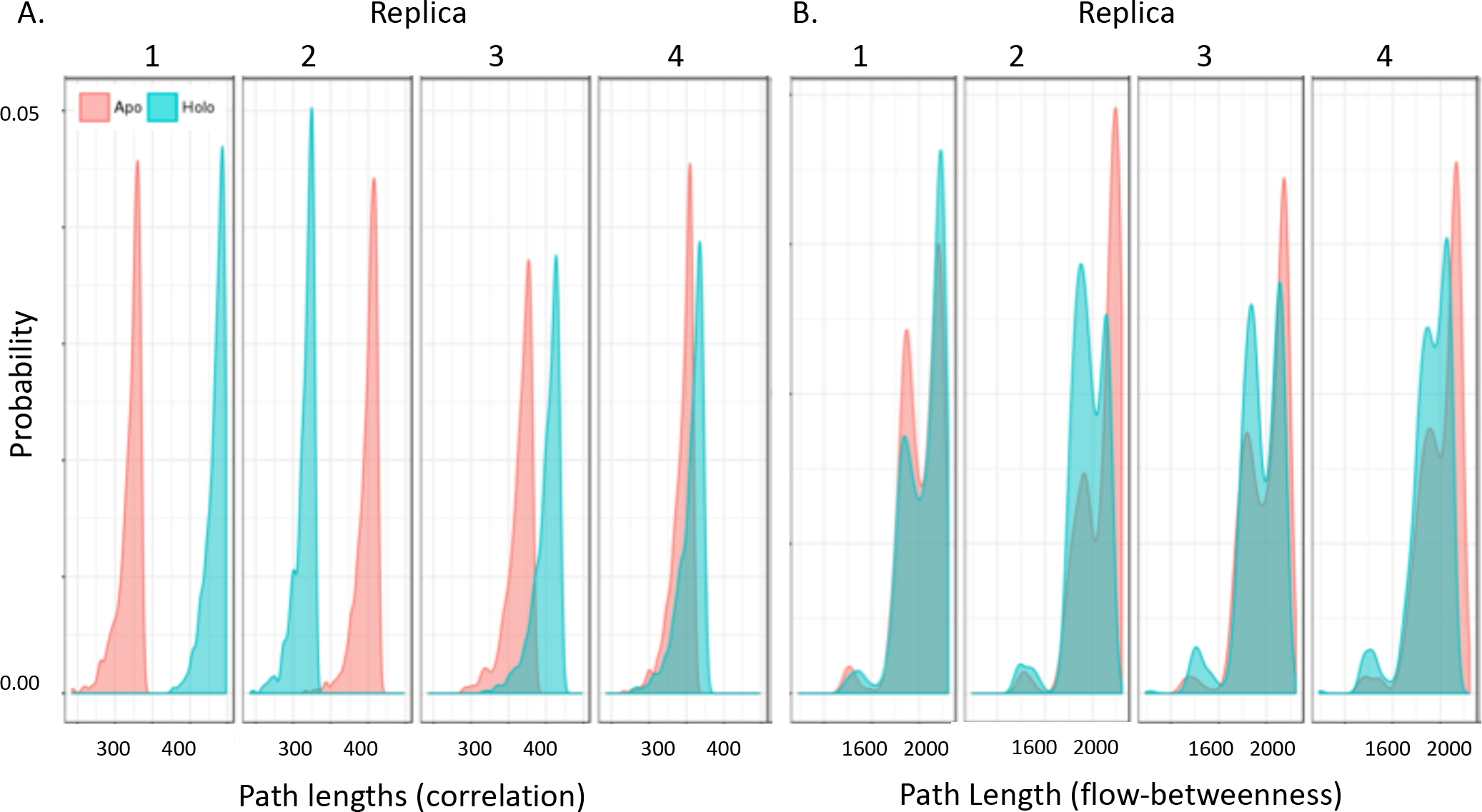
Distribution of path length. Histogram of the top 700 shortest path between source and sink nodes are plotted for apo (red) and holo (green) trajectories for each replica of 50 ns simulation. The length of the path are measured by correlation edge weighting (A) or current-flow betweenness edge weighting (B).

Perhaps the most important message brought by this analysis is the importance of multiple replicas. Although we demonstrated that the flow-betweenness edge weighting provides much more stable allosteric network topology, our analysis still shows the fluctuation between independent replicas. The importance of replicas have been pointed out for conformational sampling and 10-20 replicas are necessary to produce more reliable conclusion than a single long simulation^32-33^. Although this is not the focus of current study, using just four replicas, we emphasize that multiple replicas are also important for dynamical network analysis, even in the situation when no large conformational change is expected from the perturbation of allosteric communication.

A general point worth discussing is the length of simulation needed for conducting dynamical network analysis. It has been shown that the activation of the IGPS is in millisecond timescale^24^. Even though the time required for motion to propagate a sufficient conformational change may span milliseconds, the timescale required to transmit motion between individual amino acids in close proximity to one another is on the order of picoseconds to nanoseconds. Thus, even relatively short (in reference to the timescale of the larger conformational change) simulations could still allow one to reconstruct potential pathways along which relevant motion or energy transfer occurs. Dynamic network analysis based on correlations or pair-wise interactions is not suitable to apply directly to long-time scale trajectories that sample multiple states. This is because the time-averaged correlation or interaction is used to construct the adjacent matrix. Ideally, one would employ such long simulations in an endeavor to gather sufficient sampling to apply Markov State modeling or a similar method to determine relevant micro states. Network analysis could then be conducted over individual micro states instead of over the entire trajectory.

### Delta-network between apo and holo systems

To identify the changes between the allosteric network caused by the substrate binding, we first compared the frequency of each residue (node) that appeared in any of the top 700 paths in the apo and holo systems. Figure 6 shows all residues that have exhibited an average frequency of larger than 0.05 among the four replicas, with the frequencies from each individual replica shown as dots. Once again, the fluctuations of the node frequencies between replicas are significantly lower in the suboptimal paths generated using current-flow betweenness edge weighting (bottom) than using correlation edge weighting (top).

**Figure 6:**
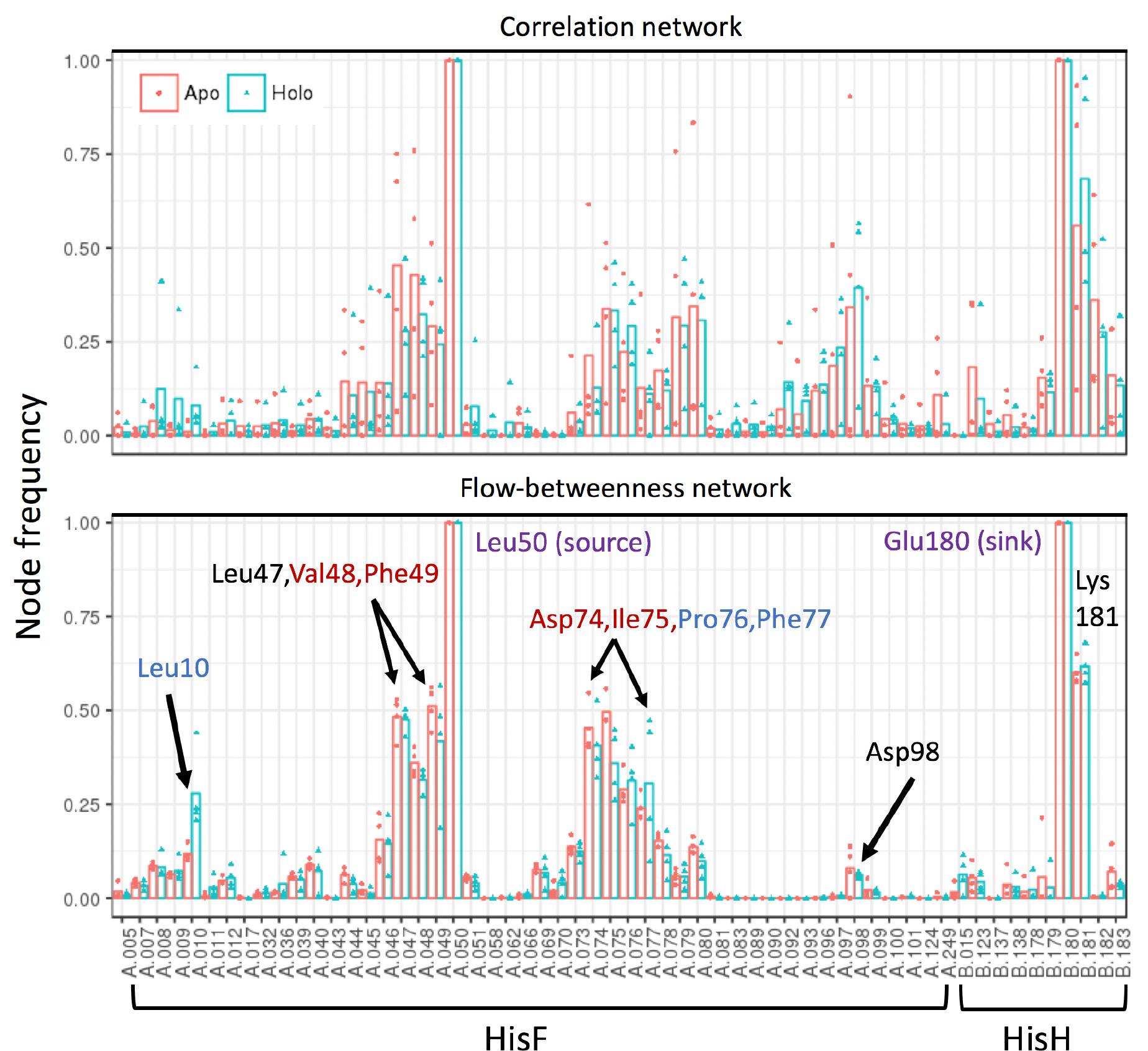
Node frequencies in the suboptimal pathways. The node frequency larger than 5% in top 700 paths are shown for (Top) suboptimal paths ranked by correlation edge weighting; (Bottom) suboptimal paths ranked by flow-betweenness edge weighting. Triangle dots depict observed frequency for individual replicas, bars indicate mean frequencies over all replicas. Residues are labeled in the following color code: blue if the node frequency is higher in the holo than in apo systems, red if the frequency is lower in holo, in black if there is no frequency change. The letter A in front of the residue ID indicates domain HisF, and letter B indicates the residues in domain HisH.

It is encouraging that among the nodes that appear more than 5% of the top 700 suboptimal paths, the current-flow betweenness captured, with much less noise, the conserved salt bridge residues K181-D98, the cation-*π* interaction residues W123-R249, as well as the majority of the conserved residues previously predicted to be involved in allosteric regulation (R5, D11, E46, P76, T78, D98, K99 in domain HisF, N15, W123, Y138, H178, E180, K181 in domain HisH).^34^ However, to compare the change between apo and holo networks rigorously, the node frequency change alone is not sufficient. This is illustrated in the following delta-network analysis. To generate a delta-network, the difference (apo minus holo) in edge frequency (*η*) are calculated from the flow-betweenness networks. The frequency difference for each edge can be loaded into VMD NetworkView plugin as a delta-edge frequency matrix in order to view the delta-network directly on the protein structure. As shown in Figure 7, the average edge frequency difference *Δη* between apo and holo are plotted on IPGS holo structure with PRFAR substrate bound. Since we define *Δη = η*(apo) *-η*(holo), the edge with positive *Δη* (shown in red bond) is more important in signal transmission between source (binding site) and sink (catalytic site) for apo state, and the edge with negative *Δη* (shown in blue bond) means this edge frequency is increased in the suboptimal path when PRFAR bound, thus becomes more important in holo state. The nodes along the delta-network are also colored based on the change in the node frequency in Figure 6 (bottom).

**Figure 7:**
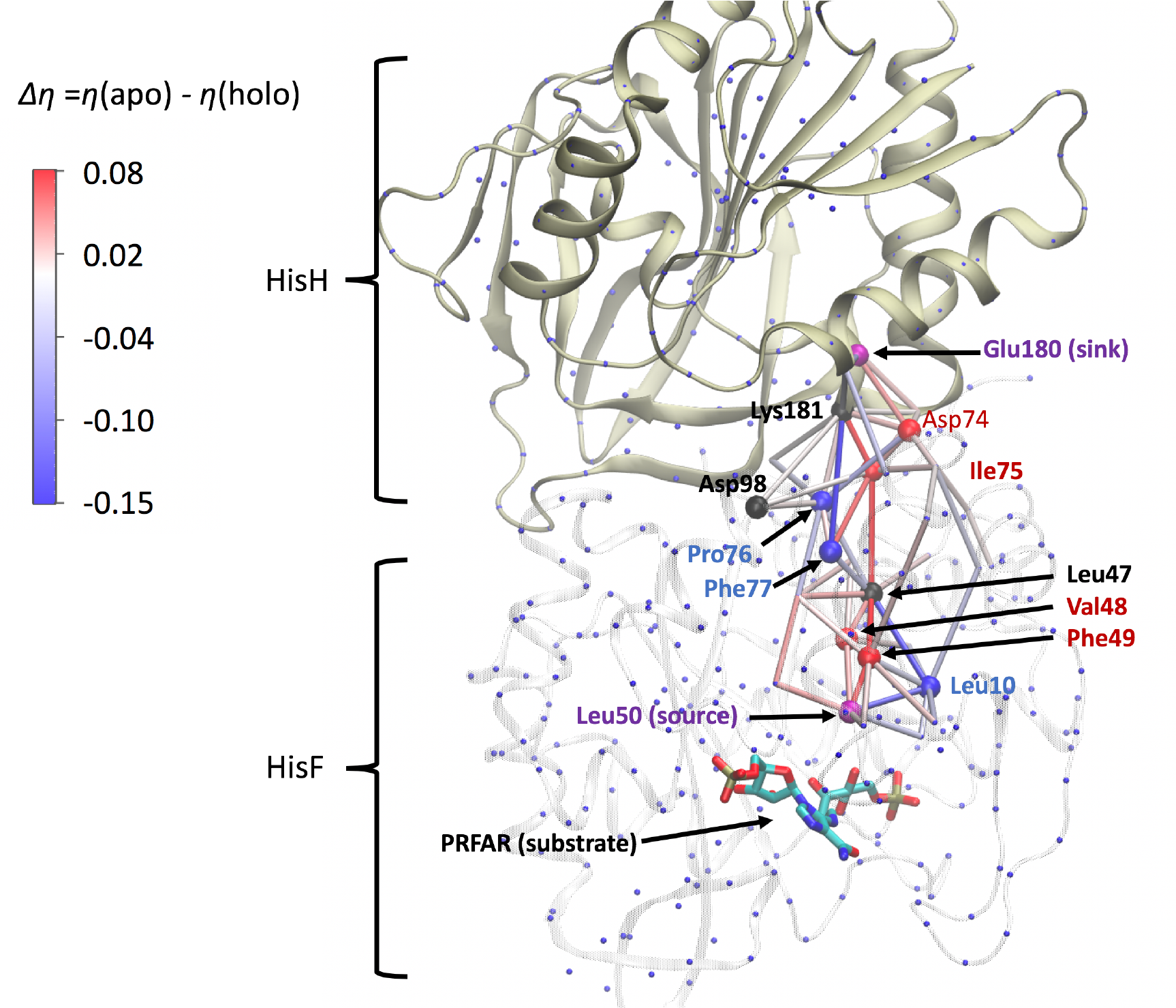
Delta-network between apo and holo IGPS. The average edge frequency difference *Δη* between apo and holo are plotted on IPGS holo structure with PRFAR substrate bound. The edge is colored with *Δη* color scale. Since *Δη **=** η*(apo) - *η*(holo), red edge (positive) is more important in apo, and blue (negative) edge is important in holo state. The important nodes are represented as balls with the color code as in Figure 6. Edges with |*Δη*| less than 0.01 are omitted here for clarity. Nodes which exhibited a decreases path usage frequency in the holo state compared to the apo state are shown as red spheres while nodes that exhibit an increase in path usage frequency in the holo state are marked with blue spheres. This color scheme matches the edge coloring as depicted in the color scale bar to the left of the figure. The source and target nodes are marked with purple spheres. Finally, the Asp98 and Lys181 residues, for which no change was observed is marked with a black spheres. The PRFAR ligand is shown using the licorice representation.

It is interesting to note that our delta-network has successfully located a key domain (L50-F49-V48-L47) associated with millisecond scale motions observed in NMR studies^35^. The other major cluster of nodes (D74-I75-P76-F77) represents a set residues at the interface between the HIF-HIS domains and is likely serving as a bridge. These domains are apparently located on a relatively flexible loop structure. We speculate that might be one of the reasons that those residues were not highlighted by the previous NMR observations aimed at identifying millisecond timescale motions, since motion of flexible regions will likely be somewhat more rapid. It is also important to note that although the Asp98-Lys181 edge was captured among the top 700 suboptimal paths, the edge usage frequency is rather low. Previous studies aimed at connecting network analysis with NMR observations of millisecond scale motions identified a pair of communication pathways leading through separate regions of the IGPS protein^24-25, 35^. The network identified in current study probably represents only a subset of those communication pathways because our particular choice of source and sink node. Glu180 is only one of the three potential catalytic triad residue C84-H178-E180. Leu50 is one of the residues lies in close proximity to the PRFAR molecule in the holo form. It is likely that Asp98-Lys181salt-bridge, while known to be critical to allosteric interaction, was not central to transmission between the specific source-target pair selected. Thus it should be noted that the set of paths presented here is by no means representative of the full communication network of IGPs. This underscores the importance of selecting an accurate subset of source and sink target residues. Nevertheless, this shortcoming in fact highlights the key feature of the flow betweenness metric, namely that it is capable of reducing contributions from edges and nodes that are not directly involved in connecting the given source and target residues, yet does not eliminate their contributions entirely.

Perhaps the most interesting point brought up by our delta-network concept is that nodes Leu47 and Lys181 have negligible change in the node frequency between apo and holo states (colored in black in Figure 6 and 7). However, the edges connecting them have significant change. This is because the information transmitting through a node is the sum of the all the edges connecting to it. For node Leu47, the edges connecting Leu47-Phe49, Leu47-Val48, Leu47-Ile75 show decreased frequency in holo state (red color), but the edges connecting Leu47-Leu10, Leu47-Phe77, Leu47-Pro76 show increased frequency in holo state (blue). The sum of all edges’s *Δη* connecting Leu47 hence cancels out, leading to no change in the Leu47 node frequency. The same situation is seen for node Lys81. Therefore, even when a node is known to contribute the transmission of the allosteric signaling, a change in the node usage frequency is not a necessary condition for the change in the allosteric network. The delta-network introduced here is thus a more rigorous and quantitative way to detect subtle network changes caused by external perturbation.

## Conclusion

Network analysis is a highly sought-after method for investigating how binding or mutation changes the protein allosteric signaling propagation, or for identifying a druggable allosteric binding site. It was shown previously that the commonly utilized methods for network construction based upon pairwise correlation can suffer from instabilities when used in assessing the relative importance of residues and residue to residue interactions in a particular allosteric signal transmission. This is because changes in the correlation between a pair of residues, regardless the type of correlation, may not necessarily imply an increase in path usage. This is particularly relevant when considering systems over which the residue-to-residue contacts and correlations fluctuate during the simulation time even a system has reached the thermodynamic equilibrium.

Inspired by electrical circuit analysis, we introduced current-flow betweenness metric that is robust in ranking how important each residue or residue-residue interaction is in propagating the allosteric signal in a protein dynamical network. Using a classic example of an allosteric enzyme IGPS, we calculated the current-flow betweenness network between a substrate binding site and a remote catalytic site more than 20Å distance apart. Through a thorough comparison with suboptimal paths generated using pairwise correlation edge weighting, we demonstrate that current-flow betweenness metric yields significantly more stable edge ranking, path length distribution, and node frequencies, regardless the choice of contact map criteria or node representations. Moreover, current-flow betweenness provides a more theoretically sound assessment of the relative importance of edges or nodes in a network since it implicitly includes contributions from all possible paths connecting a given set of nodes. Since current-flow betweenness scores exhibit less pronounced fluctuations, they can allow for identification of subtle but relevant changes in edge or node path usage with greater sensitivity than when correlation is used directly as edge weighting. Therefore, current-flow betweenness can be seen to have great potential utility for future works in which allosteric signaling pathway analysis are needed for understanding the molecular mechanism of protein function or allosteric drug design.

Very recently, a work by Negre *et. al.* reported using eigenvector centrality for identifying key amino acid residues for IGPS allostery^25^. Eigenvector centrality bears several similarities to flow betweenness. The former makes use of the eigen-decomposition of the network adjacency matrix. Specifically, the first eigenvector is utilized. Flow-betweenness, on the other hand, seeks to utilize the pseudo inverse of the matrix Laplacian, which is directly derived from, but is not equivalent to, the network adjacency matrix (Figure 2). While we here directly compute the Moore-Penrose pseudo inverse, it has also been suggested that a sufficient approximation would be the first few eigenvectors of the Laplacian matrix eigenvector decomposition^27^. Given the relation that both methods have to eigen-decomposition of similar matrix based network representations, it is likely that both methods will provide similar utility. While in this study we focus on determination of a robust allosteric network between a predefined residue pairs, how these two methods compare to each other, as well as to other betweenness and centrality metrics, both in mathematical underpinnings and their utility with respect to network analysis applications, may indeed be a good avenue for future study.

Finally, it should be noted that current-flow betweenness could be employed upon any topology and weighting scheme (provided that all edge weights are positive). For example, current-flow betweenness can be used with any correlation coefficients such as the commonly used Pearson correlation coefficients, or generalized correlation coefficients based on the mutual information concept.^26^ Furthermore, this method could potentially yield improved results when coupled with interaction energy based network construction such as in the papers by Ribeiro and Ortiz^6, 8^, which also show to increase the stability of the network over contact based topology and correlation weighting schemes. For current IGPS simulations, the contact maps are relative stable between simulation replicas. In systems that contact map stability is an issue, for instance, in larger protein or longer simulation, the linear smoothing function introduced here can mitigate artifacts that arise due to the use of arbitrary cutoff distances and frequencies when constructing contact topologies. In such case, current-flow betweenness combined with linear smooth function can overcome, to a large extent, the loose topological definitions, thus could be extremely useful given the rapid rise in the application of network analysis methods to modeling and simulation investigations.

## ACKNOWLEDGEMENT

This work was supported by NSF XSEDE research allocation MCB160119.

## References

1. Cui, Q.; Karplus, M., Allostery and cooperativity revisited. Protein Sci 2008, 17 (8), 1295–307.

2. Sethi, A.; Eargle, J.; Black, A. A.; Luthey-Schulten, Z., Dynamical networks in tRNA:protein complexes. Proc Natl Acad Sci U S A 2009, 106 (16), 6620–5.

3. Cheng, S.; Niv, M. Y., Molecular dynamics simulations and elastic network analysis of protein kinase B (Akt/PKB) inactivation. J Chem Inf Model 2010, 50 (9), 1602–10.

4. Bhattacharyya, M.; Bhat, C. R.; Vishveshwara, S., An automated approach to network features of protein structure ensembles. Protein Sci 2013, 22 (10), 1399–416.

5. Nussinov, R.; Tsai, C. J.; Ma, B., The underappreciated role of allostery in the cellular network. Annu Rev Biophys 2013, 42, 169–89.

6. Ribeiro, A. A.; Ortiz, V., Determination of signaling pathways in proteins through network theory: importance of the topology. J Chem Theory Comput 2014, 10 (4), 1762–1769.

7. Van Wart, A. T.; Durrant, J.; Votapka, L.; Amaro, R. E., Weighted implementation of suboptimal paths (WISP): an optimized algorithm and tool for dynamical network analysis. J Chem Theory Comput 2014, 10 (2), 511–517.

8. Ribeiro, A. A.; Ortiz, V., Energy propagation and network energetic coupling in proteins. J Phys Chem B 2015, 119 (5), 1835–1846.

9. Botello-Smith, W. M.; Alsamarah, A.; Chatterjee, P.; Xie, C.; Lacroix, J. J.; Hao, J.; Luo, Y., Polymodal allosteric regulation of Type 1 Serine/Threonine Kinase Receptors via a conserved electrostatic lock. PLoS Comput Biol 2017, 13 (8), e1005711.

10. Fernandez-Marino, A. I.; Harpole, T. J.; Oelstrom, K.; Delemotte, L.; Chanda, B., Gating interaction maps reveal a noncanonical electromechanical coupling mode in the Shaker K(+) channel. Nat Struct Mol Biol 2018.

11. del Sol, A.; Fujihashi, H.; Amoros, D.; Nussinov, R., Residues crucial for maintaining short paths in network communication mediate signaling in proteins. Mol Syst Biol 2006, 2, 2006 0019.

12. Romero-Rivera, A.; Garcia-Borras, M.; Osuna, S., Role of Conformational Dynamics in the Evolution of Retro-Aldolase Activity. ACS Catal 2017, 7 (12), 8524–8532.

13. Wagner, J. R.; Lee, C. T.; Durrant, J. D.; Malmstrom, R. D.; Feher, V. A.; Amaro, R. E., Emerging Computational Methods for the Rational Discovery of Allosteric Drugs. Chem Rev 2016, 116 (11), 6370–90.

14. Ribeiro, A. A.; Ortiz, V., A Chemical Perspective on Allostery. Chem Rev 2016, 116 (11), 6488–502.

15. Nussinov, R., Introduction to Protein Ensembles and Allostery. Chem Rev 2016, 116 (11), 6263–6.

16. Guo, J.; Zhou, H. X., Protein Allostery and Conformational Dynamics. Chem Rev 2016, 116 (11), 6503–15.

17. Stone, J.; Eargle, J.; Sethi, A.; Li, L.; Luthey-Schulten, Z. Tutorial: Dynamical Network Analysis; 2012.

18. Humphrey, W.; Dalke, A.; Schulten, K., VMD: visual molecular dynamics. J Mol Graphics 1996, 14 (1), 33–38.

19. Brandes, U.; Fleischer, D. In Centrality Measures Based on Current Flow, STACS, Springer: 2005; pp 533–544.

20. Oster, G.; Perelson, A.; Katchals.A, Network Thermodynamics. Nature 1971, 234 (5329), 393-+.

21. Stephenson, K.; Zelen, M., Rethinking centrality: Methods and examples. Social networks 1989, 11 (1), 1–37.

22. Newman, M. E., A measure of betweenness centrality based on random walks. Social networks 2005, 27 (1), 39–54.

23. Vanwart, A. T.; Eargle, J.; Luthey-Schulten, Z.; Amaro, R. E., Exploring residue component contributions to dynamical network models of allostery. J Chem Theory Comput 2012, 8 (8), 2949–2961.

24. Rivalta, I.; Sultan, M. M.; Lee, N. S.; Manley, G. A.; Loria, J. P.; Batista, V. S., Allosteric pathways in imidazole glycerol phosphate synthase. Proc Natl Acad Sci U S A 2012, 109 (22), E1428–36.

25. Negre, C. F. A.; Morzan, U. N.; Hendrickson, H. P.; Pal, R.; Lisi, G. P.; Loria, J. P.; Rivalta, I.; Ho, J.; Batista, V. S., Eigenvector centrality for characterization of protein allosteric pathways. Proc Natl Acad Sci U S A 2018, 115 (52), E12201–E12208.

26. Lange, O. F.; Grubmuller, H., Generalized correlation for biomolecular dynamics. Proteins 2006, 62 (4), 1053–61.

27. Bozzo, E.; Franceschet, M., Effective and efficient approximations of the generalized inverse of the graph Laplacian matrix with an application to current-flow betweenness centrality. arXiv preprint arXiv:1205.4894 2012.

28. Jo, S.; Kim, T.; Iyer, V. G.; Im, W., CHARMM-GUI: a web-based graphical user interface for CHARMM. J Comput Chem 2008, 29 (11), 1859–65.

29. Jorgensen, W. L.; Chandrasekhar, J.; Madura, J. D.; Impey, R. W.; Klein, M. L. K., Comparison of simple potential functions for simulating liquid water J Chem Phys 1983, 79 (2), 926–935.

30. D.A. Case, R. M. B., D.S. Cerutti, T.E. Cheatham, III, T.A. Darden, R.E. Duke, T.J. Giese, H. Gohlke, A.W. Goetz, N. Homeyer, S. Izadi, P. Janowski, J. Kaus, A. Kovalenko, T.S. Lee, S. LeGrand, P. Li, C. Lin, T. Luchko, R. Luo, B. Madej, D. Mermelstein, K.M. Merz, G. Monard, H. Nguyen, H.T. Nguyen, I. Omelyan, A. Onufriev, D.R. Roe, A. Roitberg, C. Sagui, C.L. Simmerling, W.M. Botello-Smith, J. Swails, R.C. Walker, J. Wang, R.M. Wolf, X. Wu, L. Xiao and P.A. Kollman AMBER 16, University of California: San Francisco, 2016.

31. Glykos, N. M., Software news and updates carma: A molecular dynamics analysis program. J Comput Chem 2006, 27 (14), 1765–1768.

32. Knapp, B.; Ospina, L.; Deane, C. M., Avoiding False Positive Conclusions in Molecular Simulation: The Importance of Replicas. J Chem Theory Comput 2018.

33. Grossfield, A.; Zuckerman, D. M., Quantifying uncertainty and sampling quality in biomolecular simulations. Annu Rep Comput Chem 2009, 5, 23–48.

34. Amaro, R. E.; Sethi, A.; Myers, R. S.; Davisson, V. J.; Luthey-Schulten, Z. A., A network of conserved interactions regulates the allosteric signal in a glutamine amidotransferase. Biochemistry 2007, 46 (8), 2156–2173.

35. Lisi, G. P.; East, K. W.; Batista, V. S.; Loria, J. P., Altering the allosteric pathway in IGPS suppresses millisecond motions and catalytic activity. Proc Natl Acad Sci U S A 2017, 201700448.

